# A Suite of Biochemical and Cell-Based Assays for the Characterization of KRAS Inhibitors and Degraders

**DOI:** 10.1101/2024.07.20.604418

**Authors:** Medhanie Kidane, Rene M. Hoffman, Jennifer K. Wolfe-Demarco, Ting-Yu Huang, Chi-Ling Teng, Luis M. Gonzalez Lira, Jennifer Lin-Jones, Gabriel Pallares, Jane E. Lamerdin, Nicole B. Servant, Chun-Yao Lee, Chao-Tsung Yang, Jean A. Bernatchez

**Affiliations:** Research and Development and Technology Transfer, Eurofins DiscoverX, LLC, 11180 Roselle Street, Suite D, San Diego, CA 92121, United States of America; Research and Development, Eurofins DiscoverX Products, LLC, 42501 Albrae Street, Fremont, CA 94538, United States of America; Eurofins Panlabs Discovery Services Taiwan, Ltd., 25 Wugong 6th Road, Wugu District, New Taipei City 24891, Taiwan

**Keywords:** KRAS, dissociation constant, target engagement, cell signaling, targeted protein degradation

## Abstract

KRAS is an important oncogenic driver which is mutated in numerous cancers. Recent advances in the selective targeting of KRAS mutants via small molecule inhibitors and targeted protein degraders have generated an increase in research activity in this area in recent years. As such, there is a need for new assay platforms to profile next generation inhibitors which improve on the potency and selectivity of existing drug candidates, while evading the emergence of resistance. Here, we describe the development of a new panel of biochemical and cell-based assays to evaluate the binding and function of known chemical entities targeting mutant KRAS. Our assay panels generated selectivity profiles and quantitative binding interaction dissociation constants for small molecules and degraders against wild type, G12C, G12D, and G12V KRAS, which were congruent with published data. These assays can be leveraged for additional mutants of interest beyond those described in this study, using both overexpressed cell-free systems and cell-based systems with endogenous protein levels.

**TABLE OF CONTENTS/ABSTRACT GRAPHIC:** 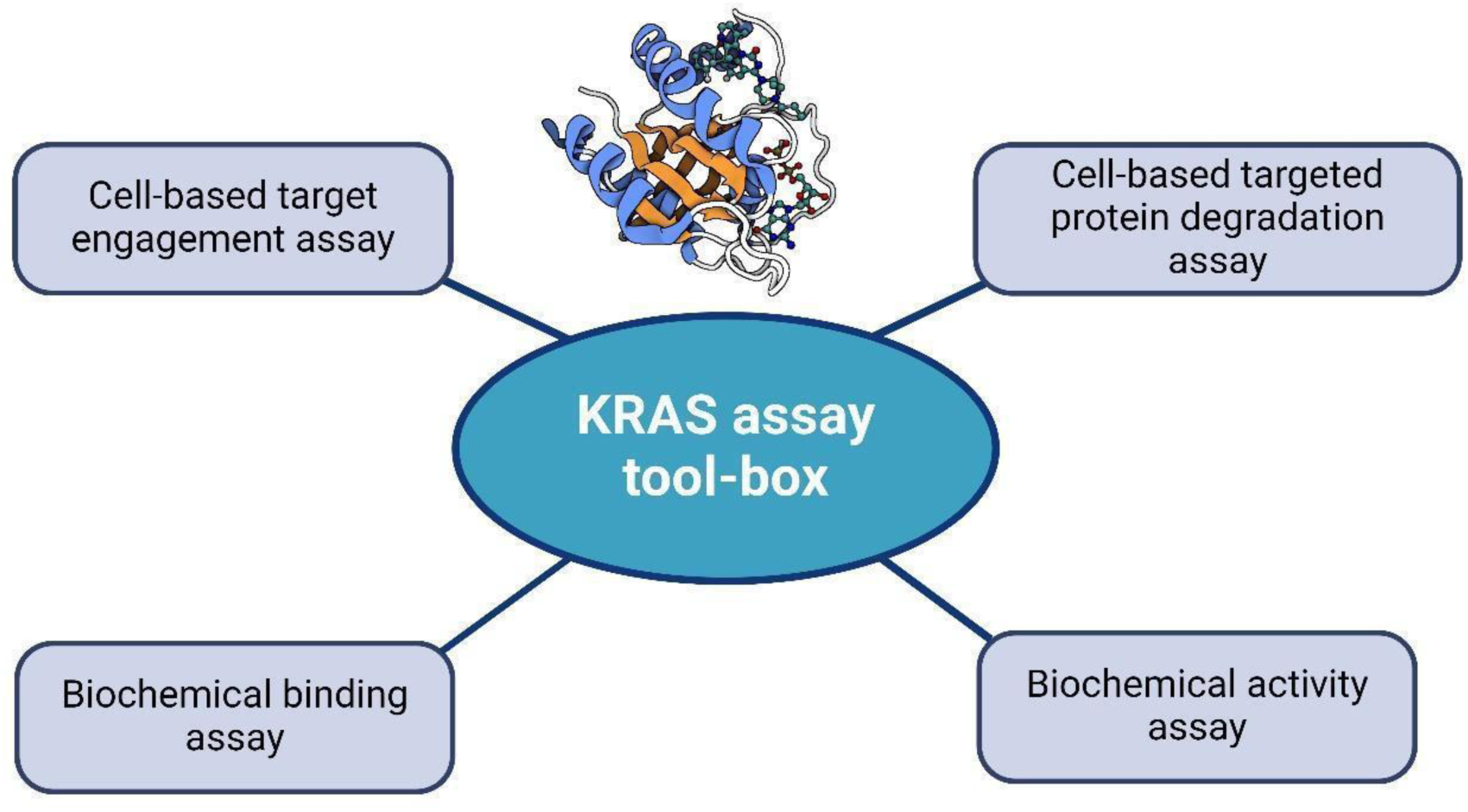

---

Kirsten rat sarcoma (KRAS) is a small GTPase that plays an important role in the regulation of biochemical signaling for cellular growth, proliferation, differentiation, and migration. Through its cycling between a guanosine triphosphate (GTP)-bound “on” state and a guanosine diphosphate (GDP)-bound “off” state, the protein can respond to extracellular signaling and propagate a biochemical signal via interaction with downstream effectors^1^. Activating mutations of KRAS are among the most common driver mutations in human cancers, and frequently alter the protein’s structure such that GTPase activating proteins (GAPs) are less able to hydrolyze GTP to GDP when bound to KRAS. This in turn biases mutant KRAS to adopt an “on” confirmation, even in the absence of an extracellular signal, thereby driving aberrant downstream cellular signaling and tumor formation^1^. A majority of KRAS mutations occur at three codon positions (G12, G13, Q61), though mutations at other positions in KRAS also can lead to the deregulation of its activity^2^. These mutations have a high occurrence in lung^3^, colorectal^4^ and pancreatic cancers^5^.

The prevalence of KRAS mutations has made the protein a prime target for oncology drug discovery for decades. However, KRAS was thought to be undruggable until the recent development of KRAS(G12C) inhibitors^6–9^, including MRTX849 (adagrasib)^10,11^ and AMG510 (sotorasib)^12,13^ that have reached the clinic. Efforts are being made to expand inhibitor space to other mutants beyond KRAS(G12C)^14–24^. Furthermore, manipulation of the ubiquitin– proteasome system to achieve targeted degradation of hyper-active or over-abundant proteins that cause diseases has become a new approach for therapeutic discovery, including for KRAS mutants^25^.

The recent clinical success of the KRAS(G12C) inhibitors MRTX849^10^ and AMG510^12^ has ignited extensive interest to adapt this modality into bi-functional degraders that could reduce hyperactive KRAS mutant proteins for cancer therapeutics. Commercially available LC2 is a mutant selective KRAS Degrader that consists of MRTX849 chemically linked to a ligand for the von Hippel Lindau (VHL) E3 ligase^25,10,26^. It has been reported that LC2 induces selective degradation of KRAS(G12C) protein with a DC_50_ around 0.25-0.76 μM without causing degradation of other forms of KRAS proteins in cancer cell lines. The degradation of the KRAS(G12C) protein was shown to lead to a decrease of MAPK signaling in these cells and consequently growth suppression^25,27^.

Accurate measurement of the dissociation constants (K_D_s) for KRAS inhibitors such as the KRAS G12D-selective inhibitor MRTX1133^15,28,29^ has sometimes been challenging, particularly when the affinity of the ligand for the target is very high and the K_D_ for the interaction is below the detection limit of the technology used for the measurement^15^. K_D_ values for high-affinity ligands have been successfully determined in the past for kinases^30,31,32,33^ using an assay system that tags the kinase with a DNA probe that can detect picomolar quantities of protein via qPCR. This allows for the accurate determination of the true thermodynamic dissociation constant for a ligand without hitting the tight-binding limit of the assay for picomolar affinity ligands^33^.

Traditional assays for protein degradation and turnover, such as cell viability assays, ELISAs and Western blots, are laborious and time-consuming^34–37^. Hence, a homogenous cell-based assay without the need for reliable antibodies is desirable for the discovery of new protein degraders in a high throughput manner. Despite the common use in drug discovery of cell lines that overexpress a target protein, these systems sometimes have limitations when accurately determining the potencies of many degraders.

To better characterize KRAS-targeting compounds for drug discovery efforts in this space, we developed both biochemical and cell-based assays to assess target engagement by small molecules, and functional assays to assess the effect of KRAS degraders on endogenous protein levels in relevant cancer cell models.

The biochemical competition-based binding assay described in this paper for KRAS is based on the qPCR-amplicon tagging technology mentioned above. In this assay, a DNA-tagged protein of interest is collected as part of a mammalian cell protein extract, and then incubated in the presence of a capture ligand immobilized on magnetic beads. The KRAS-binding warhead of the immobilized capture ligand is taken from a compound series that has been shown to engage the switch II binding pocket of KRAS, which has been a successful strategy for the allosteric inhibition of this target’s activity^15^. The afore-mentioned components are further incubated in the presence of a dimethylsulfoxide (DMSO) vehicle control or a test competitor compound. If the test compound can compete with the immobilized capture ligand for binding to the target protein, less protein is enriched on the bead in this case than in the presence of a vehicle control. After quantification of the protein that was captured on the bead via qPCR, a lower qPCR signal would be observed for a compound that was able to compete with the immobilized capture ligand for binding to the target protein.

The nucleotide exchange assay (NEA) is a widely used technique to study RAS functional activity between the “off” (GDP) and “on” (GTP) states of the protein; this process is mediated by guanine nucleotide exchange factors (e.g., SOS1)^38,39^. To complement the inhibitor data collected from the developed biochemical binding assays, we performed activity assays where fluorophore-labeled GDP was used and GDP to GTP exchange across KRAS(wild-type, WT), KRAS(G12C), KRAS(G12D), and KRAS(G12V) was directly measured through time-resolved fluorescence energy transfer (TR-FRET)^40^, as previously described^39^.

To meet the need for homogenous cell-based degradation assays, a new type of cell line was developed based on the enzyme fragment complementation (EFC) method to capture endogenous expression of target proteins. EFC is a homogeneous, robust and scalable detection assay system that enables the measurement and ranking of ligand potencies, discovery of the mechanism of action (MOA) of a compound, the performance of functional screens, and identification of novel compounds that engage the target^41–44^. This approach is based on two recombinant β-galactosidase (β-gal) enzyme fragments that act as an enzyme acceptor (EA) and an enzyme donor (ED). Separately, the fragments are inactive, but when combined, they form an active β-gal enzyme that hydrolyzes its substrate to produce a chemiluminescence signal^41^. The ED fragment is relatively small (42 amino acids) and can be easily knocked into an endogenous target locus in the cell model of choice to create an ED-target fusion protein that is expressed at physiologically relevant levels. In addition, the ED fragment contains no lysines, which are residues that can become covalently linked to ubiquitin^41,45^. Thus, no proteins are artificially degraded due to the introduction of the ED fragment. The endogenous expression levels of ED-target fusion proteins more accurately reflect physiologically relevant protein levels than overexpression systems and thus allow for a more biologically relevant potency assessment of protein degraders.

In this study, biochemical competition binding assays were performed using KRAS(WT), KRAS(G12C), KRAS(G12D) and KRAS(G12V) NFκB DNA-binding domain fusions tagged with qPCR amplicons to detect the protein in the assay readout. We were able to quantitatively measure K_D_ values for MRTX1133 and other KRAS-targeting compounds across the constructs tested and were able to recapitulate published rank order selectivity for these small molecules.

In addition, three KRAS cell lines (KRAS(G12C), KRAS(G12D) and KRAS(WT)) were generated by knocking in an ED fragment (and the relevant mutation in KRAS) into the *KRAS* locus by CRISPR technology. We demonstrated that these cell lines are excellent assay tools for investigating protein degraders and inhibitors. First, we showed their utility in functional testing of protein degradation by demonstrating that the LC2 PROTAC (proteolysis-targeting chimera) selectively induces protein degradation in KRAS(G12C) cells and not in KRAS(WT) and KRAS(G12D) cells, matching the published selectivity of the compound for KRAS(G12C)^25^. LC2 has been shown to selectively bind KRAS(G12C) through its MRTX849 warhead^10^ and to recruit the E3 ligase VHL via a VH032 ligand^26^ to ubiquitinate and ultimately degrade KRAS(G12C)^25^. Second, we adapted a target engagement workflow ^46,47^ for these cell lines and demonstrated that they are useful to interrogate the direct binding between KRAS mutants and their specific inhibitors and gained rank order information for the affinity of these inhibitors to their targets.

Together, these assays provide a framework for the interrogation of the binding and functional parameters of small molecule inhibitors and targeted protein degraders of KRAS and its mutants and can be applied to the characterization of new chemical matter being developed against mutant forms of the protein.

## RESULTS

### Biochemical Binding Assays to Determine Quantitative Affinities of KRAS Switch II Pocket Binders

To provide a new platform to study binding of small molecules to the switch II binding pocket of WT and mutant KRAS proteins, we developed a set of biochemical competitive binding assays, based on a technology platform that has been previously applied to study ligand binding to kinases^30,31,32,33^. KRAS(WT), KRAS(G12C), KRAS(G12D) and KRAS(G12V) constructs with N-terminal fusions of the DNA binding domain of NFκB^48^ were transiently transfected in HEK293 cells. Protein extracts containing these constructs were harvested, then mixed with a DNA probe that anneals with the NFκB fusion domain of the constructs and which contains a qPCR amplicon for protein detection. Finally, the protein extracts were incubated with magnetic beads baited with the biotin-small molecule conjugate compound **1**, which contains a ligand moiety previously described as a switch II pocket binder (**Figure 1A**)^15^. Incubation of the extract with the baited beads was conducted in the presence of either escalating concentrations of a switch II pocket binder (the competitor compounds MRTX1133^15,28,29^, MRTX849^10,11^ and AMG 510^12,13^) (**Figure 1B**) or a DMSO vehicle control. Following washing and elution of the residual protein bound to the baited beads, a qPCR reaction was conducted to quantify the protein in the eluate. A scheme showing the binding assay principle is depicted in **Figure 2**.

**Figure 1.**
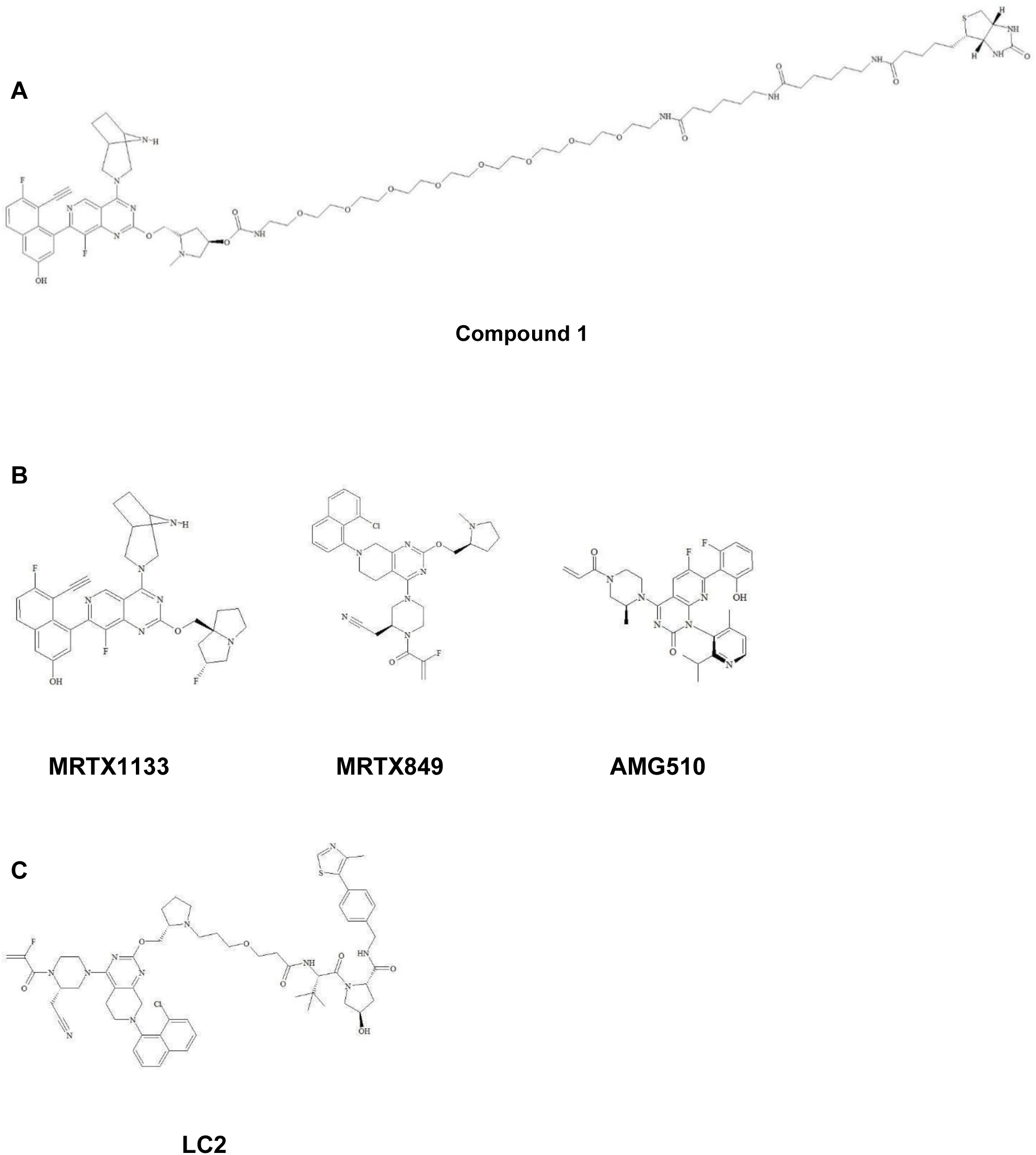
Chemical Structures of Compounds Used in This Study. **(A)** Biotin-conjugated capture ligand for the competitive binding assay. **(B)** KRAS switch II pocket-binding small molecule inhibitors. **(C)** KRAS heterobifunctional degrader.

**Figure 2.**
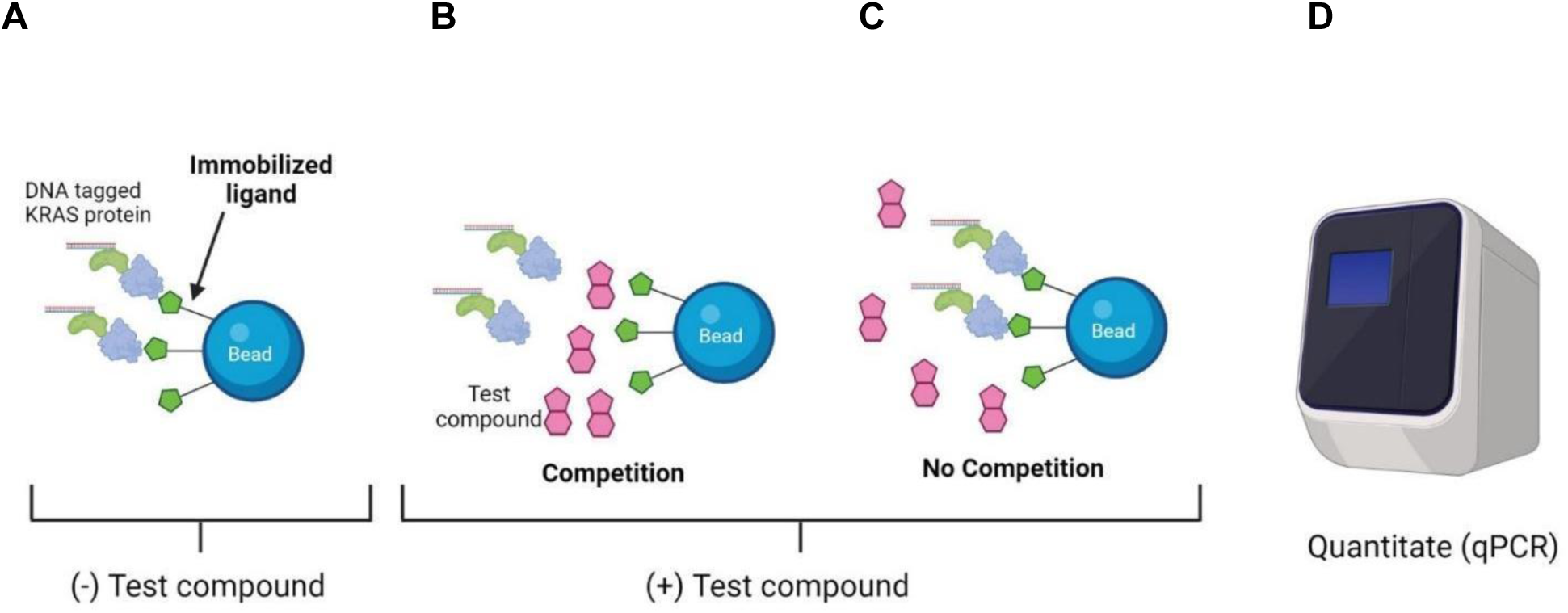
Overview of the Biochemical Competition Binding Assay Principle. **(A)** Engagement of the DNA tagged KRAS protein with the immobilized ligand on the bead (bait molecule) in the absence of a test compound and the presence of the DMSO vehicle control is shown. Following washing of the beads and elution of KRAS protein captured on the beads, a high protein concentration is expected in the eluate. **(B)** In the presence of a test compound that competes with the bait molecule for binding to KRAS, a reduced amount of target protein is captured on the bead. Following washing of the beads and elution of residual KRAS protein, the expected protein concentration in the eluate is lower. **(C)** In the presence of a test compound that does not compete with the bait molecule for binding to KRAS, a high amount of target protein is captured on the bead. Following washing of the beads and elution of residual KRAS protein, the expected protein concentration in the eluate is high. **(D)** At the end of the assay, a qPCR reaction is conducted to quantify the amount of amplicon tagged KRAS protein present in the eluate.

Curve fitting of the competitor dose response data yielded thermodynamic dissociation constants for the competitor compounds (sample curves for MRTX1133 affinity are shown in **Figure 3**), as described in the Methods. Importantly, the bait loading on the beads was adjusted so that the K_D_(app) ≈ K_D_ for the competitor molecules tested, as has been previously described for this type of assay^33^. The obtained K_D_ values for MRTX1133, MRTX849 and AMG510 are shown in **Table 1**.

**Figure 3.**
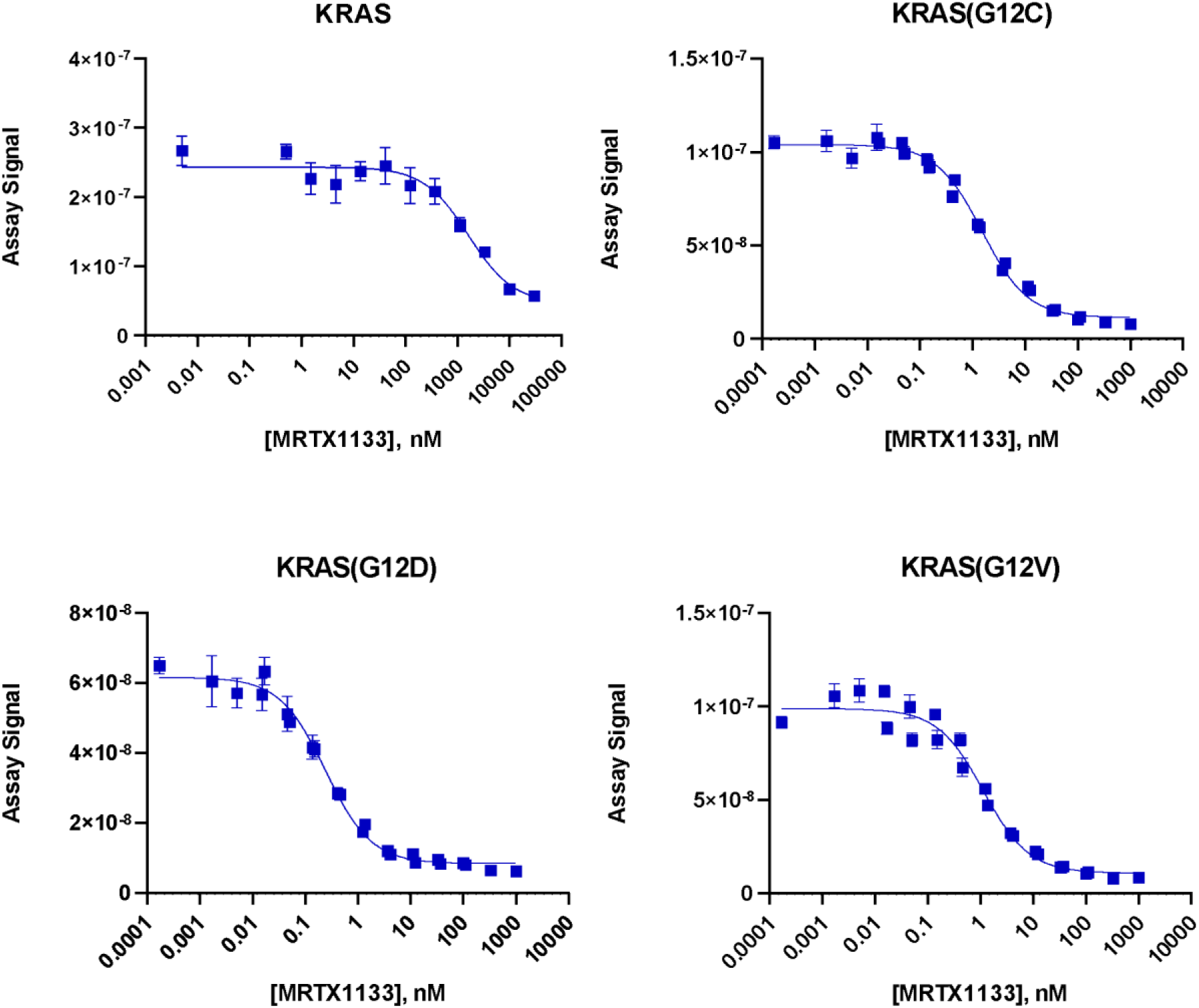
Sample Binding Curves for MRTX1133 Engagement with WT and mutant KRAS in the Developed Biochemical Binding Assay. Shown are representative curves for the fitting of qPCR quantification data for recovered KRAS protein in the assay eluate for escalating concentrations of MRTX1133. Error bars on data points represent the standard deviation, n=4 for KRAS(WT), n=8 for KRAS(G12C), KRAS(G12D) and KRAS(G12V).

**Table 1.** Thermodynamic Dissociation Constants Measured for KRAS Inhibitors Targeting the Switch II Binding Pocket. Average K_D_ values for MRTX1133, MRTX849 and AMG510 binding to WT and mutant KRAS are shown for replicates collected over three independent experiments ± standard deviation.

| Protein | Binding Incubation Time | Compound Name | Average $K_D$ (nM) | Number of Replicates |
| --- | --- | --- | --- | --- |
| KRAS | 24hr | MRTX1133 | $2560 \pm 56$ | 6 |
|  |  | MRTX849 | >20,000 | 6 |
|  |  | AMG510 | >20,000 | 6 |
| KRAS(G12C) | 1hr | MRTX1133 | $2.35 \pm 0.10$ | 12 |
| | | MRTX849 | $9.59 \pm 2.09$ | 6 |
| | | AMG510 | $220 \pm 47$ | 6 |
| KRAS(G12D) | 1hr | MRTX1133 | $0.40 \pm 0.11$ | 12 |
|  |  | MRTX849 | >20,000 | 6 |
|  |  | AMG510 | >20,000 | 6 |
| KRAS(G12V) | 1hr | MRTX1133 | $1.72 \pm 0.11$ | 12 |
|  |  | MRTX849 | >20,000 | 6 |
|  |  | AMG510 | >20,000 | 6 |

We measured a K_D_ of 400 pM for MRTX1133, a selective KRAS(G12D) non-covalent inhibitor, against the G12D mutant; this is in contrast with the previously estimated K_D_ of the compound of ∼0.2 pM, where the predicted K_D_ of the compound was below the accuracy limit of the SPR instrument used to measure the dissociation constant^15^. The measured K_D_ values for this compound for the G12C and G12V mutants was higher than the value measured for the G12D mutant (2.35 nM and 1.72 nM, respectively). As expected, the measured affinity of MRTX1133 in this study was lower for KRAS(WT) (K_D_ of 2560 nM). Together, these results highlight the selectivity of MRTX1133 for the G12D mutant over the other two G12 mutants tested and the WT protein. In addition, we provide quantitative K_D_ values for MRTX1133 for each of these proteins, with a measured subnanomolar K_D_ value for KRAS G12D.

The covalent switch II pocket-binding inhibitor MRTX849^10^ displayed a high level of binding selectivity based on our data: we obtained a K_D_ of 9.59 nM for KRAS(G12C), and no binding was observed for MRTX849 against the KRAS WT, G12D and G12V protein at compound concentrations up to 20 µM. As with MRTX849, we also observed very selective binding of AMG510^12^ for the G12C mutant, with a K_D_ of 220 nM for this protein, and no binding of AMG510 against WT, G12D and G12V proteins at concentrations up to 20 µM.

### Biochemical Activity Assay

To confirm the affinity rank order profile for MRTX1133 and AMG510 that we observed in our developed biochemical binding assays, TR-FRET-based activity profiling of these compounds against KRAS(WT), KRAS(G12C), KRAS(G12D) and KRAS(G12V) was performed (**Figure 4**). IC_50_ values for the inhibition of nucleotide exchange are shown in **Table 2**. Matching the compound affinity trend seen in the biochemical binding assays, MRTX1133 selectively inhibited KRAS(G12D) (IC_50_ = 0.14 nM) over KRAS(WT) (IC_50_ = 5.37 nM), KRAS(G12C) (IC_50_ = 4.91 nM) and KRAS(G12V) (IC_50_ = 7.64 nM). In addition, AMG510 inhibited KRAS(G12C) (IC_50_ = 8.88 nM), but no inhibition of KRAS(WT), KRAS(G12D) and KRAS(G12V) was observed up to 100 µM of compound; these results matched the relative affinity profile obtained for AMG510 in the biochemical binding assays.

**Figure 4.**
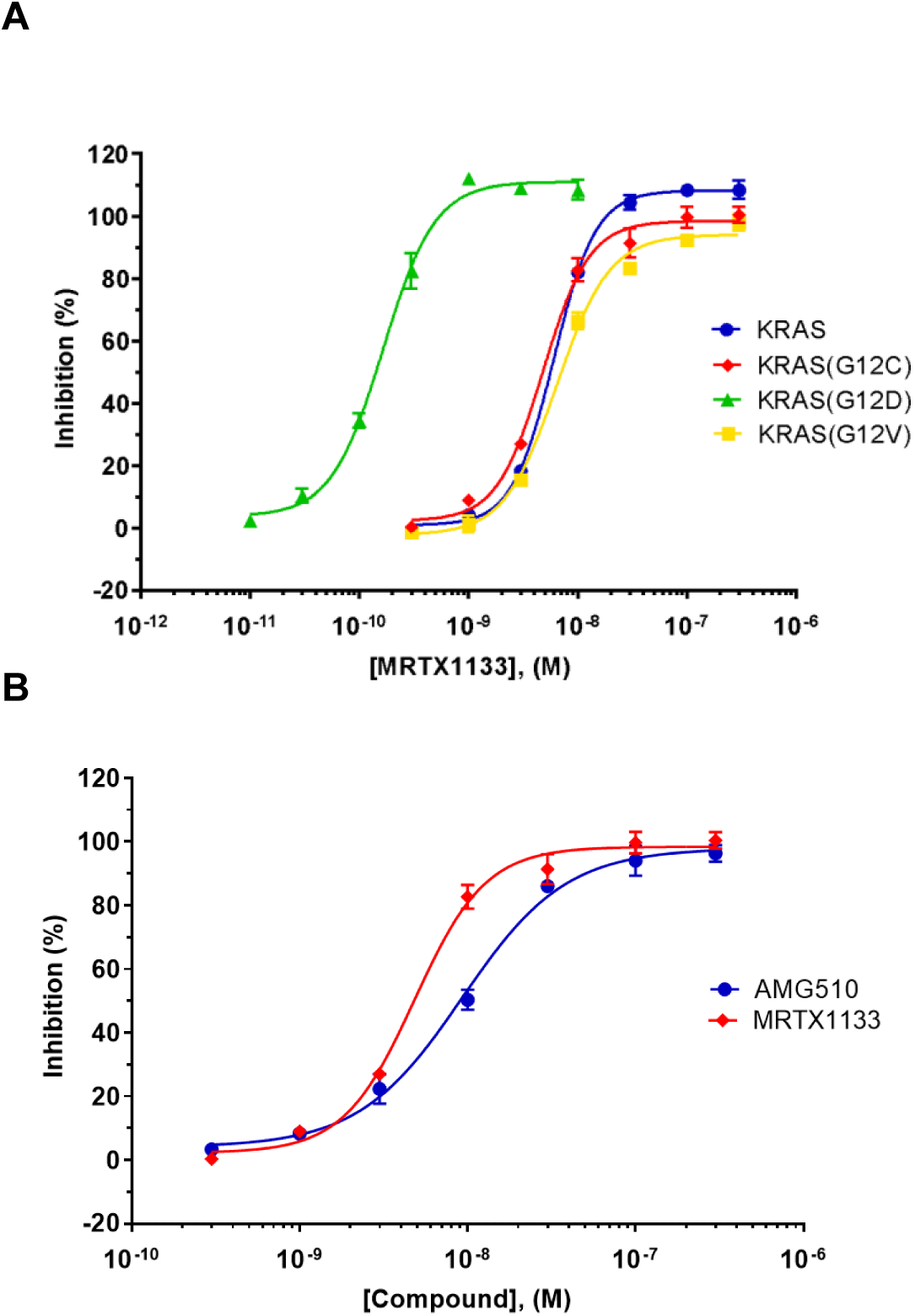
The Inhibitory Effects of MRTX1133 and AMG510 on Nucleotide Exchange Against KRAS(WT) and Mutants. **(A)** Relative potencies for MRTX1133 against KRAS(WT), KRAS(G12C), KRAS(G12D) and KRAS(G12V). **(B)** Inhibition potencies for MRTX1133 and AMG510 in the KRAS(G12C) assay. Error bars on data points represent the standard error of the mean for three independent experiments.

**Table 2.**
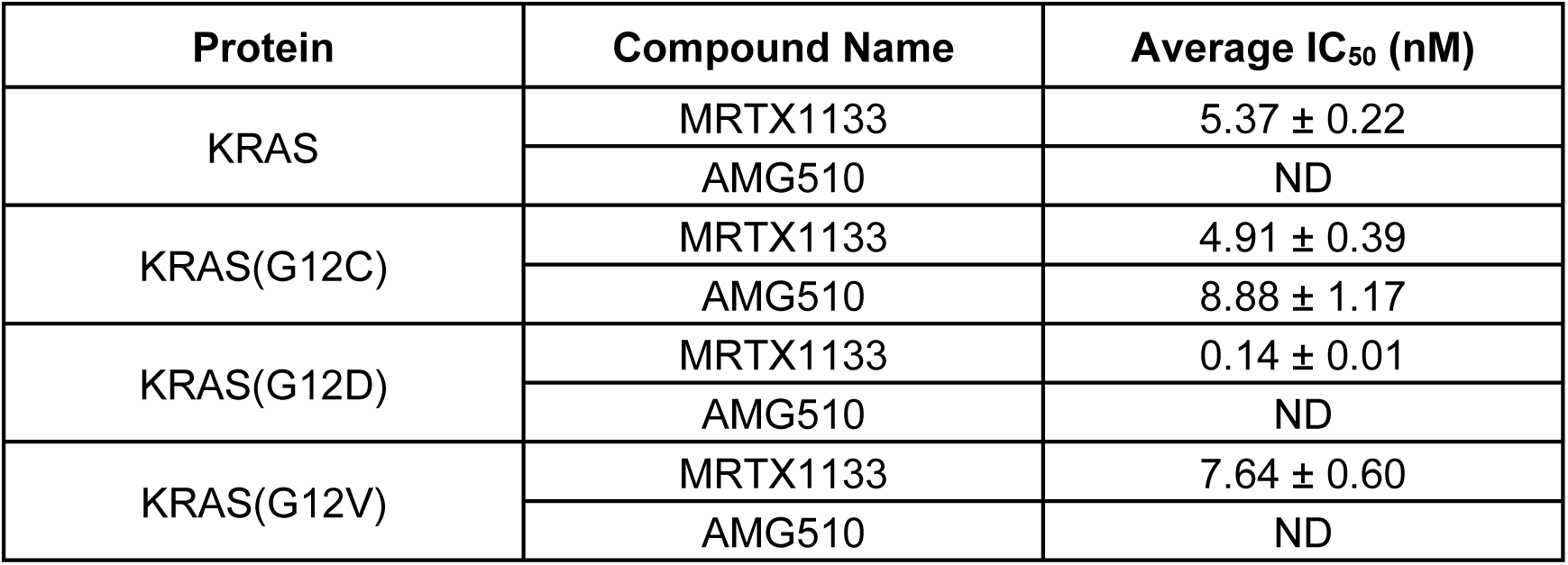
IC_50_ Values of MRTX1133 and AMG510 against KRAS(WT) and Mutants. Average IC_50_ values are shown for replicates collected from three independent experiments ± standard error of the mean. ND: No inhibition detected at 100 μM compound.

### CRISPR-Mediated Knock-In

To capture the endogenous expression of KRAS in an A549 cell model, we introduced an ED tag (ePL or enhanced ProLink) into the *KRAS* locus to create an N-terminal tagged ED-KRAS fusion protein via the CRISPR-Cas9 system (**Figure 5**). The editing efficiencies of gRNAs that recognize sequences near the translation start site on *KRAS* exon 2 were first evaluated. gRNA (CR11.1) with an editing rate above 70% was used along with double-stranded DNA (dsDNA) donors that contained an ED tag flanked by homologous sequences with a G12C mutation or G12 to correct *KRAS^G12S^*to *WT* in the A549 cells. The events of homologous recombination were detected by PCR-based molecular analysis and the occurrence of in-frame knock-in (KI) were confirmed by EFC activities. Cell pools underwent limiting dilution to isolate single-cell derived clones that harbored homozygous ED-*KRAS* alleles. We estimated that the homologous recombination frequencies were around 15-20%. Individual clones were further evaluated for their performance in functional assays, namely targeted protein degradation and target engagement. Best performing clones were tested for functional passage stability up to 15 passages and clones showing stable assay windows were used in the study. The KRAS(G12D) cell line was generated by introducing a G12D mutation into the established KRAS(G12C) line. Therefore, these two lines share virtually identical genetic backgrounds.

**Figure 5.**
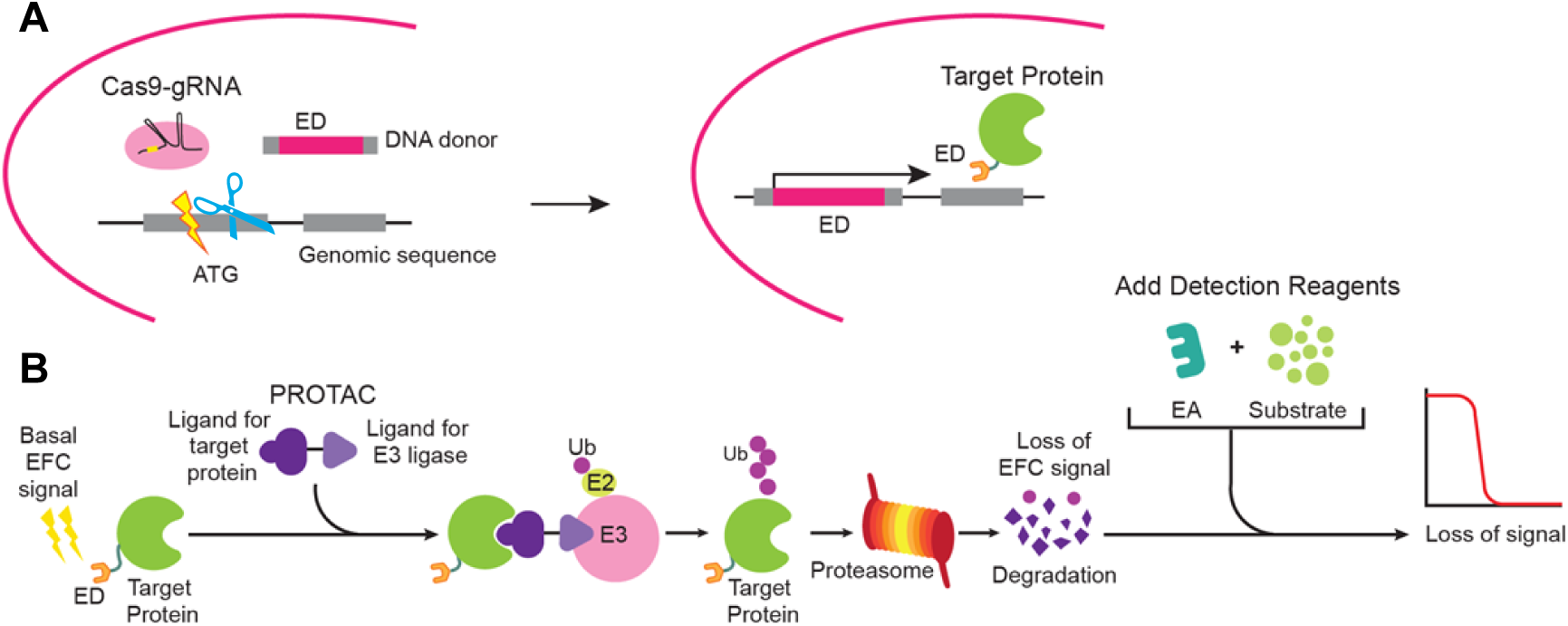
Schematic Presentation of CRISPR-Cas9-Mediated Knock-In and the Developed Protein Degradation Assay using Enzyme Fragment Complementation (EFC). **(A)** A KRAS cell line was generated by introducing a ribonucleoprotein complex containing gRNA and Cas9 protein along with a dsDNA donor comprised of the ED tag flanked by homologous recombination sequences into cells. Events leading to the in-frame knock-in needed to generate ED-target fusion proteins expressed at endogenous levels in a disease-relevant cell model are shown. **(B)** ED-target fusion proteins expressed at high or medium levels (basal EFC signal) are brought into proximity with an E3 ligase via a bi-functional molecule (such as a PROTAC). Subsequently, the target protein is ubiquitinated and degraded by the proteasome, resulting in a loss of EFC signal.

### Targeted Protein Degradation Assay

We tested the KRAS(G12C)-specific protein degrader LC2 (**Figure 1C**) with our three KRAS cell lines in the developed protein degradation assay. The assay was set up in homogenous 384-well format with an 11-point dose response curve, with a top dose of 10 μM and each dose tested in quadruplicate. Cells were incubated with LC2 for 18 hrs and at the end of incubation, detection reagents containing the complementing EA fragment were added to each well for chemiluminescence quantitation. Shown in **Figure 6A**, LC2 induced a loss of signal (an indication of protein degradation) in KRAS(G12C) cells with an IC_50_ of 1.9 μM, close to the previously described value^25^. In addition, protein degradation was only induced in KRAS(G12C) but not in KRAS(WT) and KRAS(G12D) cells (**Figure 6A-C)**. This finding is consistent with reports that MRTX849 is specific to the KRAS(G12C) protein^10^.

**Figure 6.**
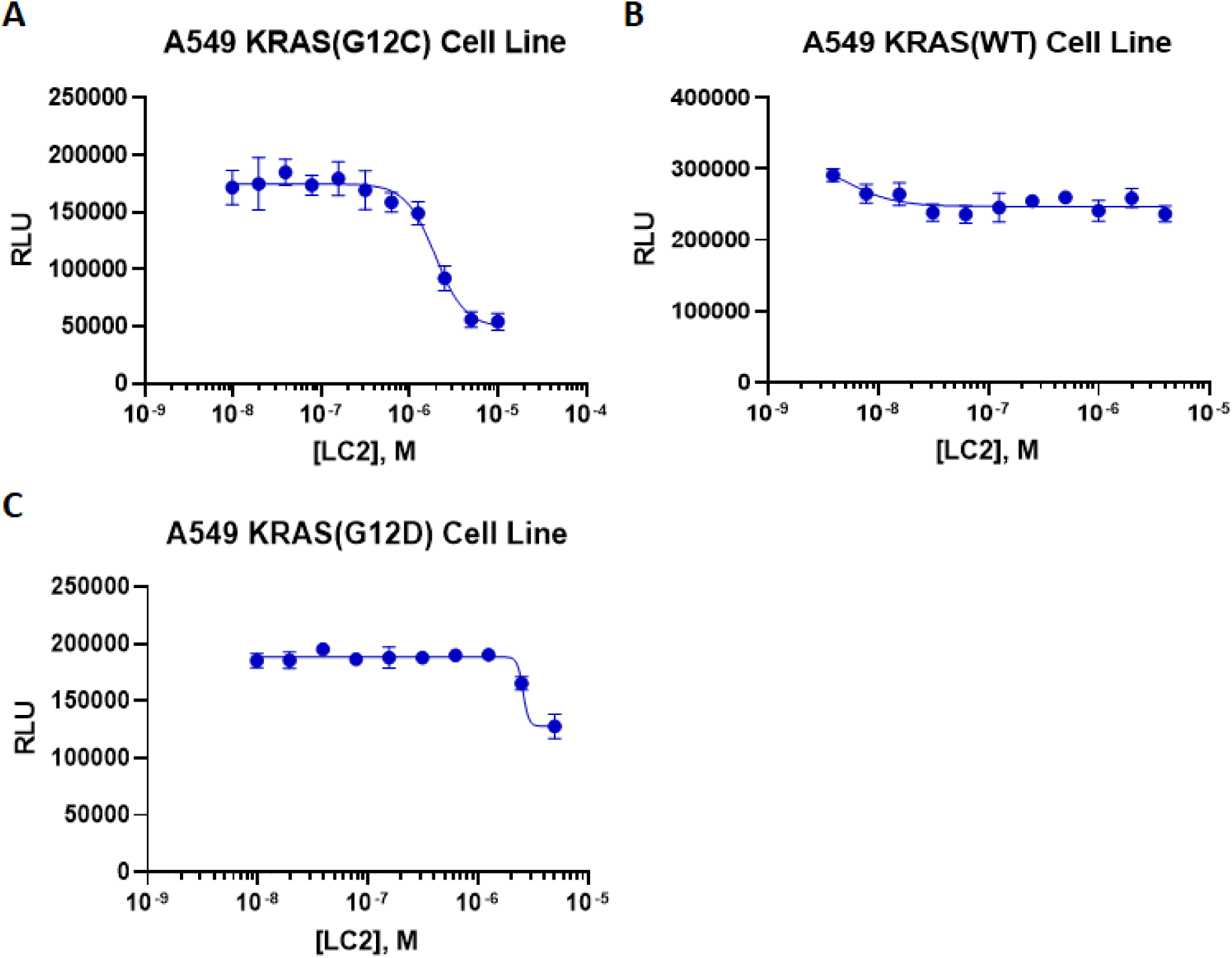
The PROTAC LC2 Selectively Induces KRAS(G12C) Protein Degradation. Three KRAS cell lines (KRAS(G12C) **(A)**, KRAS(WT) **(B)** and KRAS(G12D) **(C)**), were incubated with PROTAC LC2 for 18 hr. LC2 selectively induces KRAS(G12C) protein degradation with an IC_50_ of 1.9 µM, yet spares the other KRAS proteins tested. Error bars on data points represent the standard deviation, n>3 for KRAS(G12C), n=3 for KRAS(WT) and KRAS(G12D). RLU = relative luminescence units.

### Pulse Target Engagement Assay

Next, we explored the application of KRAS cell lines for a pulse target engagement (TE) assay to profile compounds or identify new ligands. The TE assay is based on the observation that the direct interaction between a compound and its cellular protein target increases the stability of the protein and reduces its turnover^46,47,49^. Given that most endogenous proteins have a relatively long half-life, introduction of a thermal pulse has been demonstrated to induce protein turnover, increasing the success rate of TE assay development^13,50^. Therefore, in our pulse TE assay, cells are incubated with a compound and then subjected to an elevated temperature to induce protein denaturation. Compound binding protects the target protein from denaturation, and consequently preserves enzyme complementation activities with the cellular ED-tagged fusion protein, producing a robust chemiluminescent signal. In the absence of compound binding, the target protein forms denatured aggregates that leads to a loss of complementation ability and decreased chemiluminescent signal (**Figure 7**).

**Figure 7.**
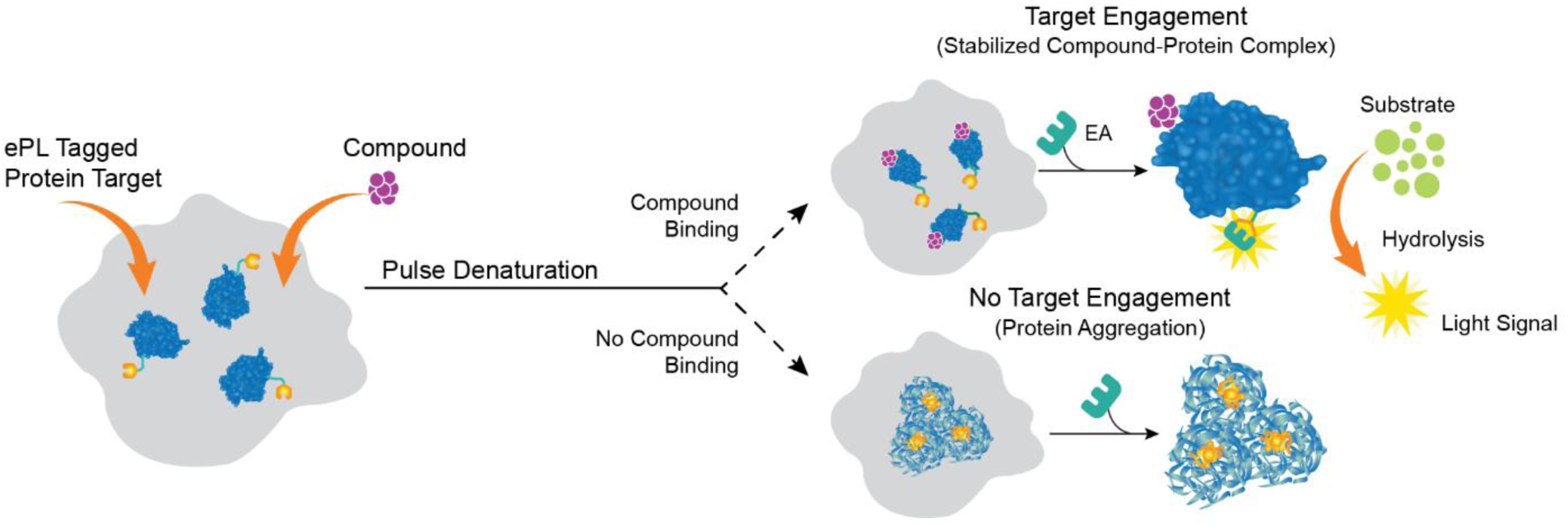
Graphical Representation of the Pulse Target Engagement Assay. The assumption made for this target engagement assay is that a direct interaction between the target protein and a compound stabilizes the protein and reduces its turnover. Cells are incubated with a compound and then subjected to a pulse of an elevated temperature. Compound-bound target protein forms a heat-stable complex. Consequently, this preserves the EFC activity in the assay and increases the chemiluminescent signal. In the absence of compound binding, the protein is denatured and aggregates and loses enzyme complementation ability, resulting in a low chemiluminescent signal.

First, a thermal shift assay was performed to determine the optimal pulse temperature for thermal insult in the pulse TE assay^46^. KRAS(G12C) cells were treated with 10 and 5 μM of MRTX849 or vehicle control (1% DMSO) for 1 hr, followed by a 3 min incubation at a gradient of elevated temperatures. Shown in **Figure 8A**, the KRAS protein appeared relatively heat-stable with a constant chemiluminescent signal detected up to a temperature of 60°C. A thermal shift was observed at temperatures above 63°C; in the presence of MRTX849, EFC activities remained steady while samples without the compound (vehicle control) had a significant drop in EFC activities. This finding indicates a direct interaction between KRAS(G12C) protein and MRTX849 as previously described, and this interaction protects the KRAS protein from thermal denaturation. We chose 67°C as the pulse temperature since a roughly 50% difference in EFC activity was detected at this temperature between MRTX849-treated samples and the vehicle control, which would therefore provide a robust assay window.

**Figure 8.**
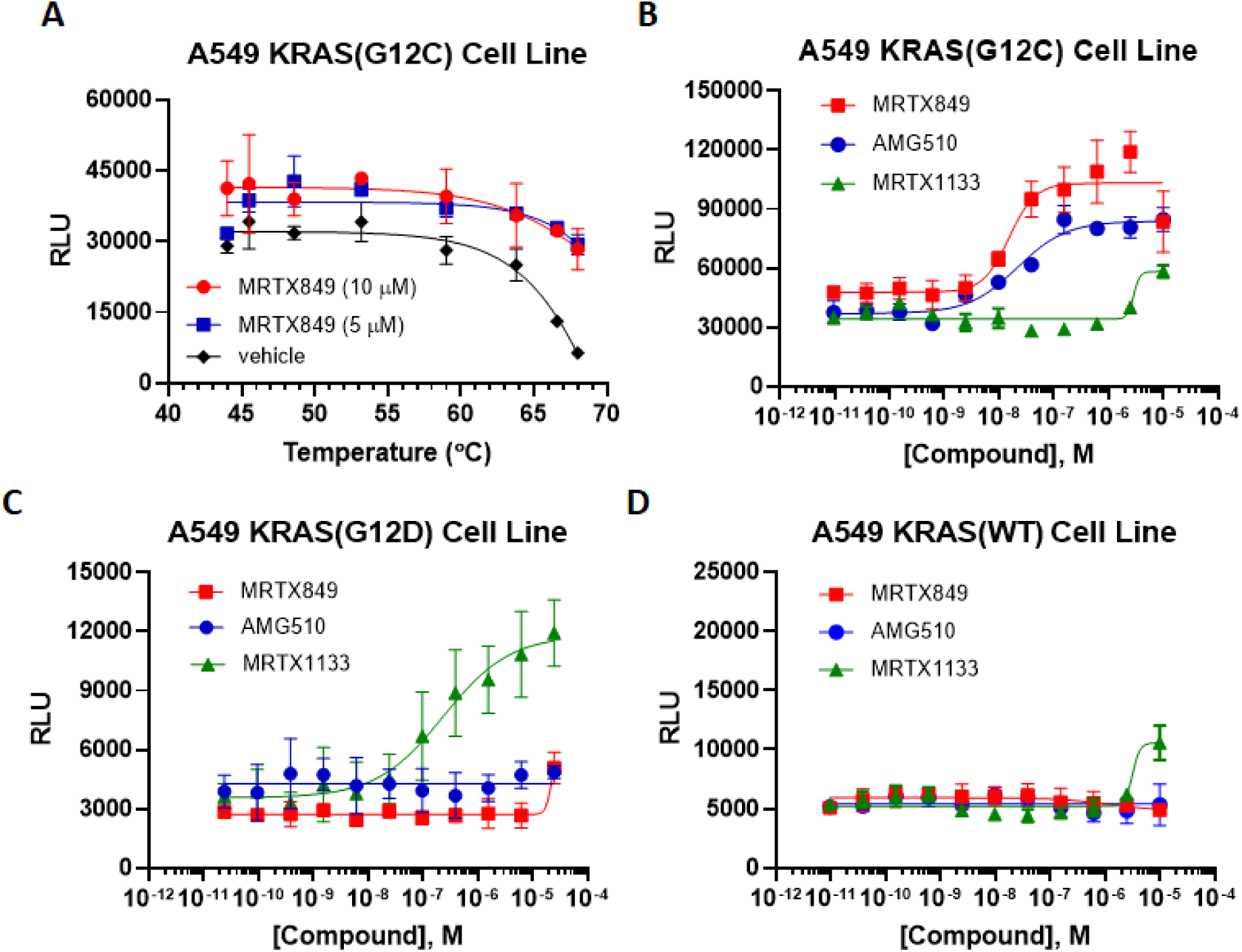
The Developed Target Engagement Assay Reveals Inhibitor Selectivity to WT and Mutant KRAS Proteins. KRAS cell lines were used in target engagement assays to profile a panel of compounds including mutant specific inhibitors. **(A)** A thermal shift assay using the KRAS(G12C) cell line and its specific inhibitor MRTX849 was first performed to determine the pulse temperature for the target engagement assay. Cells were incubated at two different concentrations of MRTX849 or vehicle (1% DMSO) for 1 hr, followed by a 3 min incubation at a gradient of elevated temperatures from 44 to 68°C. A decrease in EFC activity appeared at temperatures above 60°C and a substantial difference in EFC activity between MRTX849-treated and vehicle control-treated cells became most apparent around 66-68°C. 67°C was chosen as the pulse temperature for the target engagement assay. **(B-D)** For the target engagement assay, cells were incubated with compounds for 1 hr followed by a 3 min incubation at 67°C. MRTX849 and AMG510 showed high affinity to KRAS(G12C) protein **(B).** While MRTX1133 appears to have high selectivity for KRAS(G12D) protein **(C),** it also shows some interaction with KRAS(G12C) and KRAS(WT) protein **(D)**. Error bars on data points represent the standard deviation, n=5 for KRAS(G12C), n=4 for KRAS(WT), n=3 for KRAS(G12D).

We next performed pulse TE assays with a panel of KRAS inhibitors on three KRAS cell lines. MRTX849^10^ and AMG510^12^ are two inhibitors specific to the G12C mutant. MRTX1133 is a newly developed inhibitor that specifically targets the G12D mutant^15^. Cells were incubated with the compounds for 1 hr, followed by a 3 min incubation at 67°C. Consistent with previous reports, we observed that MRTX849 and AMG510 appeared to have high affinity only for the KRAS(G12C) protein but not for KRAS(WT) or KRAS(G12D) proteins^10,12^ (**Figure 8B-D**).

MRTX1133 showed high affinity for the KRAS(G12D) protein as previously described and some binding to KRAS(G12C) and KRAS(WT) proteins at high concentration^15^. These findings confirm that these KRAS cell lines are useful tools for compound profiling and could be used to screen new chemical matter.

## DISCUSSION

Major strides have been made in the development of KRAS-mutant selective inhibitors in cancer, most notably for KRAS(G12C) and KRAS(G12D). Unfortunately, resistance to KRAS inhibitor monotherapy has been reported in the clinic^51^; the basis for this resistance appears to be multifactorial: further mutation of KRAS which reduces inhibitor affinity for the target, changes in gene copy number, pathway rescue and alternative pathway utilization through mutation, transcriptional and epigenetic reprogramming and tumor microenvironment changes have all been proposed as mechanisms by which resistance is developed to this class of inhibitors^52^. Combination therapy^53,54,55,56^ and next-generation inhibitors that evade resistance are being proposed as solutions to this issue. As such, development of next-generation KRAS inhibitors which evade resistance mechanisms or have a higher cellular fitness cost for the development of resistance are of importance to this field. Furthermore, the development of assay platforms that can identify selective inhibitors for mutants outside of G12C and G12D (for which multiple inhibitors have already been developed) will lead to a more tailored, personalized medicine approach for patients harboring other KRAS mutations prevalent in various cancers. Recent exciting advances in pan-KRAS mutant inhibitors, such as BI-2865^22^, offer the potential for even broadly applicable treatment strategies for various mutant forms of KRAS. With the development of the new series of assays described in this paper, we have expanded the array of tools available for the development of next-generation, mutant-selective inhibitors of KRAS that circumvent existing resistance pathways. The present set of assays is amenable to expansion, including for additional KRAS mutants other than the G12C, G12D and G12V mutants described here.

We quantitatively determined the binding affinity constants for MRTX1133, AMG510 and MRTX849 across WT and G12C, G12D and G12V KRAS in biochemical assays in this work. The measured affinity constant for MRTX1133 was in the picomolar range for the G12D mutant protein we tested, consistent with the estimated high binding affinity reported previously^15^. In addition, we demonstrated that MRTX1133 bound mutant KRAS with higher affinity than for WT protein, as reported for this compound^15^. We also observed selective binding of AMG510 and MRTX849 to KRAS(G12C), as published for these compounds^12,10^. The binding selectivity profiles for MRTX1133 and AMG510 were also consistent with the data we collected for these compounds in our nucleotide exchange assays. Since the biochemical assays have been validated with the afore-mentioned reference compounds, they can be used to interrogate novel chemical matter for binding to the switch II binding pocket of KRAS for the discovery of next-generation KRAS inhibitors.

Using our developed targeted protein degradation assay, we show that the PROTAC LC2 is capable of selectively degrading KRAS(G12C) in cells while sparing KRAS(WT) and showing low levels of KRAS(G12D) degradation only at higher concentrations of the degrader. The selectivity of this compound for KRAS(G12C) recapitulates published results^25^. This assay can be used to profile novel PROTACs or molecular glues for mutant KRAS for the development of next-generation therapeutics for the treatment of mutant KRAS tumors.

In the cell-based target engagement assays, we were also able to reproduce published mutant-specific binding selectivity in our target engagement assays for three of the inhibitors tested in the biochemical system: MRTX1133 was selective for KRAS(G12D), and AMG510 and MRTX849 were selective for KRAS(G12C)^15,12,10^. These assays measure the effectiveness of the tested compounds in cells and provide important information for new drug testing that is complementary to the biochemical assays, such as evaluating compound diffusion through the cell membrane and engagement of compounds with the target expressed at physiological levels within cells.

In this work, we presented a cell-based platform for therapeutic discovery targeting KRAS mutants prevalent among many human cancers^2,3,4,5^. We demonstrated that the KRAS(G12C) cell line is highly useful for the development of assays for other KRAS mutants: we were able to simply introduce the KRAS(G12D) mutation using CRISPR KI in the cell line described herein. This allows for direct comparison of profiling results for our different mutants using a virtually identical genetic background.

In sum, we have presented a collection of biochemical and cell-based assays to study KRAS inhibitors and degraders. These assays can be used as part of a pipeline to identify and characterize the parameters for target binding or target degradation across WT and mutant KRAS proteins of novel chemical entities.

## METHODS

### Small Molecules and Nucleic Acids

MRTX1133, MRTX849 and AMG510 were purchased from MedChemExpress (Monmouth Junction, NJ) and Selleck Chemicals LLC (Houston, TX). The selective KRAS degrader LC2 was purchased from Bio-Techne Corporation (Minneapolis, MN). The biotinylated affinity probe (compound **1**), which was used for the KRAS competition binding assays, was custom synthesized by SAI Life Sciences (Watertown, MA). DMSO was purchased from Sigma-Aldrich (St. Louis, MO). The DNA probe containing the PCR amplicon used to tag the NFκB fusion domain was custom synthesized by Thermo Fisher Scientific (Waltham, MA).

### Protein Constructs and Protein Expression

Wild-type (WT) KRAS and G12C, G12D and G12V mutants (M1/R164, using NCBI entry NP_203524.1 as a reference sequence) were designed as N-terminal fusions with the DNA binding domain of NFκB (consisting of residues 35-36 fused to residues 41-359 (as described previously^48^), using UniProt entry P19838 as a reference). KRAS proteins were expressed via transient transfection in HEK293 cells. Protein extracts were harvested in M-PER extraction buffer (Pierce Biotechnology, Rockford, IL) supplemented with 150 mM NaCl, 10 mM DTT, Protease Inhibitor Cocktail Complete (Roche Diagnostics GmbH, Mannheim, Germany) and Phosphatase Inhibitor Cocktail Set II (Merck KGaA, Darmstadt, Germany) following the manufacturer’s guidelines.

### Cell Culture

Native A549 and SPRINTer^TM^ A549 KRAS WT, G12C and G12D cell lines from Eurofins DiscoverX (Fremont, CA) were maintained in DMEM medium supplemented with 10% fetal bovine serum and 1X PSG from Thermo Fisher Scientific (Waltham, MA). Cultures were maintained with a routine of medium renewal 2 to 3 times per week. Freezing medium for cryopreservation is 95% complete growth medium with 5% DMSO.

### Competition Binding Assays

Competition binding assays were designed and developed as previously described for kinases^30,31,32,33^.

Preparation of liganded beads was performed as follows: the biotinylated affinity ligand (Compound **1**) was incubated with streptavidin-coated magnetic beads (Thermo Fisher Scientific, Waltham, MA) for 30 min at 25°C. In order to remove the unbound affinity ligand and to reduce nonspecific binding of proteins in the cell lysate, the liganded beads were then blocked with excess biotin (125 nM) and washed with a blocking buffer containing SeaBlock (Pierce Biotechnology), 1% BSA and 0.05% Tween 20.

The binding reactions were prepared with the DNA tagged KRAS protein extract, beads loaded with the affinity ligand and the competitor test compounds in a binding buffer (1x PBS, 0.05% Tween 20, 10 mM DTT, 0.1% BSA, 2 mg/ml sheared salmon sperm DNA) in deep well, natural polypropylene 384-well plates, catalog number 784201 (Greiner Bio-One, Kremsmünster, Austria) in a final volume of 19.7 µL. No enzyme purification steps were performed on the protein extracts before adding them to the reaction mixture, and the protein extracts were diluted 10,000-fold in the final reaction mixture (the final DNA-tagged enzyme concentration was less than 0.1 nM). Binding assay mixtures were incubated at 25 °C with shaking for 24 h (WT KRAS assay) or 1 h (G12C, G12D and G12V KRAS assays). A 24-hour incubation time for the WT assay was required to have appreciable competition of the test compound with the bait molecule so that a usable assay window could be obtained. After the incubation period, the affinity beads were extensively treated with a wash buffer (1x PBS, 0.05% Tween 20) to remove unbound protein from the protein lysate. Using an elution buffer (1x PBS, 0.05% Tween 20 and either 20 μM MRTX1133 (WT KRAS assay) or 1 μM MRTX 1133 (G12C, G12D and G12V KRAS assays)), the beads were resuspended and incubated at 25 °C while shaking for a 30 min period. The concentration of WT or mutant KRAS in the eluates was then determined by quantitative PCR. K_D_ values for each competitor compound were determined using eleven serial threefold dilutions. For each assay, the affinity ligand concentration present on the magnetic beads was optimized to ensure that the true thermodynamic K_D_ values for competitor molecules were measured, as described in detail previously^33^.

### Data Analysis for Competitive Binding Assays

Binding constants (K_D_s) for each experiment were calculated using a standard dose-response curve fitting of the data using the Hill equation:

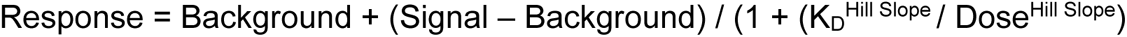

The Hill Slope was set to −1. A non-linear least square fit using the Levenberg-Marquardt algorithm was employed for curve fitting.

### Nucleotide Exchange Assays

Biotinylated human recombinant KRAS WT, G12C, G12D, and G12V proteins (Amid Bioscience, Santa Clara, CA) were pretreated with Terbium-labeled streptavidin (Tb-SA) (PerkinElmer, Waltham, MA) and BODIPY-FL-GDP (Thermo Fisher Scientific, Waltham, MA) overnight at room temperature in assay buffer containing 20 mM HEPES, pH 7.5, 50 mM NaCl, 10 mM MgCl2, and 0.01% (w/v) Tween 20. MRTX1133 and AMG510 were half-log diluted in duplicate and acoustically dispensed into 384-well plates (Corning, Corning, NY) by Echo 650 (Beckman Coulter, Brea, CA). Test articles were preincubated with pretreated KRAS/BODIPY-FL-GDP/Tb-SA for 1 h at 25°C. To initiate the reaction, pre-mixed SOS1 (Amid Bioscience, Santa Clara, CA) and unlabeled GTP (Sigma-Aldrich, St. Louis, MO) were added (total 20.2 µL per well) and incubated for 30 min at 25°C. Final concentrations of Tb-SA, SOS1, and GTP were 0.17 nM, 50 nM, and 10 μM for all assays. Final concentrations of KRAS proteins and BODIPY-FL-GDP were 10 nM and 100 nM for the KRAS WT, G12C, and G12V assays, and were 1 nM and 40 nM for the KRAS G12D assay. TR-FRET signals were obtained via excitation (337 nm) and emission (Tb: 486 nm and BODIPY: 515 nm) wavelengths using Infinite M1000 Pro spectrometer (TECAN, Männedorf, Switzerland). The inhibition percentages (%) for seven doses were analyzed and IC_50_ values were determined using GraphPad Prism.

### CRISPR-Mediated Knock-In and KRAS Cell Line Generation

The gRNAs (crRNA), HiFi Cas9 Nuclease V3, and ATTO^TM^ 550 tracrRNA used in this study to generate ED-tagged KRAS cell lines were purchased from Integrated DNA Technologies (Coralville, IA). Double-stranded DNAs containing ED or ePL (enhanced ProLink) were purchased from Eurofins Genomics Blue Heron (Bothell, WA). The gRNAs (crRNA) were designed using CRISPick, a design tool developed by the Broad Institute, with a CRISPRko mechanism by SpyoCas9 enzyme (*S. pyogenes*, NGG) on the Human Reference Genome GRCh38 (https://portals.broadinstitute.org/gppx/crispick/public). A gRNA (CR11.1, GAATATAAACTTGTGGTTGT) was used to generate KRAS(G12C) and KRAS(WT) cell lines. The gRNA was first annealed with fluorescence-labeled (ATTO-550) tracrRNA in a 1:1 ratio to form an RNA duplex, followed by incubation with Cas9 nuclease to generate Cas9-gRNA ribonucleoprotein (RNP). Cas9-gRNA RNP and double-stranded DNA (dsDNA) donors that contained ED fragments flanked by 120-180 bps of homologous sequences and the WT G12 sequence or G12C mutation were delivered into native A549 cells using the Neon Electroporation System from Thermo Fisher Scientific (Waltham, MA). At 16 hours post-electroporation, cells with high fluorescence at 550 nm were sorted to enrich the genome-edited cell populations. PCR-based molecular analysis was performed to verify homologous recombination events. These cell pools underwent limiting dilution to generate single cell derived clones. Homozygous KI clones were identified by sequencing analysis and clones with the best assay windows for the protein degradation assay or the target engagement assay were chosen for this study. gRNA CR11.2 (GTAGTTGGAGCTGGTGGCGT) and a single-stranded DNA donor containing the G12D mutation were introduced into the KRAS(G12C) cell line to generate the KRAS(G12D) cell line following a similar protocol to the one described above.

### Targeted Protein Degradation Assay

Using the ED-tagged KRAS WT, G12C or G12D cell lines and the EFC detection kit (PathHunter® ProLabel®/ProLink™ Detection Kit, Eurofins DiscoverX, Fremont, CA), protein degradation assays were set up to test the potency of PROTAC LC2 in 384-well format in 11-point dose response (quadruplicate wells/dose), with the last well in each dose curve containing the vehicle control. Cells were detached from culture dishes with accutase (Innovative Cell Technologies, San Diego, CA) and collected in conical tubes. Spent medium and accutase were removed by centrifugation and the cells were resuspended in an appropriate amount of AssayComplete^TM^ Cell Plating 0 Reagent (CP0, Eurofins DiscoverX, Fremont, CA) to allow a cell seeding of 5,000 cells in 20 µL per well in a 384-well assay plate (white, clear flat bottom, Corning, Corning, NY). LC2 was diluted in CP0 to 50 µM (5X concentration for a top dose of 10 µM in the final assay) followed by 2-fold serial dilutions to generate the remainder of the 10 doses in the dose curve. 5 µL of the 5X stock of LC2 was added to each well and 5 µL of CP0 was added to the vehicle control wells. The assay plates were placed in an incubator at 37°C with 5% CO_2_ for 18 hr. Working detection reagent solution was prepared by mixing a 4:1:1 ratio of the ED (PL/PK) detection reagent as described in the manufacturer’s user manual. 30 µL of the working detection reagent solution was added to each well at the end of compound incubation and the plates were left in the dark at room temperature for 1h. Plates were read using an Envision plate reader (Perkin Elmer, Shelton, CT) using a 0.1 sec / well integration time. The dose-response curves were generated using GraphPad Prism (GraphPad Software, Boston, MA).

### Pulse Target Engagement Assay

This workflow was adapted from the InCell Pulse Target Engagement Assay (Eurofins DiscoverX, Fremont, CA). A thermal shift assay was first performed to determine the optimal temperature for thermal insult in the pulse target engagement assay. KRAS(G12C) cells were collected and diluted in complete medium that allowed for a cell seeding of 5,000 cells in 40 µL per well in an 8-point dose curve (triplicate wells per dose) in 96-well PCR plates (black, Thermo Fisher Scientific, Waltham, MA). Working stocks (5X) of MRTX849 (50 and 25 μM) were made in complete medium with 5% DMSO. 10 μL of MRTX849 working stock (5X) was added to each well to generate final concentrations of 10 and 5 μM with 1% DMSO. For vehicle control samples, 10 μL of 5% DMSO in complete medium was added to each well. The plates were placed in an incubator at 37°C, 5% CO_2_ for 1 hr, followed by a 3 min incubation at a gradient of elevated temperatures between 44 and 68°C in a C1000 Touch Thermal Cycler (BioRad Life Science, Hercules, CA). The plates were then cooled at 25°C for 3 mins. 60 µL of the working detection reagent solution (ProLabel/ProLinK Detection Kit, Eurofins DiscoverX, Fremont, CA) described above was added to each well at the end of the cool down. The plates were left in the dark at room temperature for 1h before reading chemiluminescence signal on an EnVision plate reader.

The pulse target engagement assay was set up in a 96-well format with an 11-point dose response curve (triplicates/dose) and vehicle control. Cells were collected and diluted in complete medium that allowed for a cell seeding of 5,000 cells in 40 µL per well in 96-well PCR plates (black, Thermo Fisher Scientific, Waltham, MA). Compounds were diluted in complete medium to 125 µM (5X concentration for a top dose of 25 µM in 5% DMSO) followed by a 4-fold serial dilution to make the rest of the 10 doses. 10 µL of 5X stocks of serially diluted compounds were added to each well in the assay plate and 10 µL of complete medium was added to the vehicle control well. The assay plates were placed in an incubator at 37°C, 5% CO_2_ for 1 or 5 hours. The plates were placed in a thermal cycler for a 3 min incubation at 67°C, followed by another 3 min incubation at 25°C to cool down. 60 µL of the working detection reagent solution (same as above) was added to each well and the plates were left in the dark at room temperature for 1h before reading chemiluminescence signal using an EnVision plate reader.

## ACKNOWLEDGEMENTS

The authors acknowledge members of Eurofins DiscoverX, LLC and Eurofins DiscoverX Products, LLC for their help with initial discussions surrounding the project and for feedback on the manuscript. Figures for the Table of Contents graphic and for the assay schemes were generated using BioRender.com. Curve fitting for dose response figures was performed using GraphPad Prism (GraphPad Software, Boston, MA).

## AUTHOR CONTRIBUTIONS

M.K. developed the biochemical competitive binding assays, contributed to the generation of the dissociation constant data, the design of the clones and the design of experimental approaches, and analyzed the biochemical binding assay data. R.M.H. generated KRAS cell lines and performed experiments for the target engagement and targeted protein degradation cell-based assays. J.K.W.D. contributed to the generation of the dissociation constant data, the design of experimental approaches and analyzed the biochemical binding assay data. C.T.Y. designed the CRIPSR-mediated knock-in scheme. T.Y.H. and C.L.T. performed the nucleotide exchange assays, contributed to the design of experimental procedures and the data analysis. L.M.G.L. contributed to the clone design for the biochemical binding assays. J.L.J., G.P., J.E.L., N.B.S. and C.Y.L. contributed to the data analysis. C.T.Y. and J.A.B. wrote the manuscript, led the design of the cell-based and biochemical binding assays, respectively, and analyzed the data. All authors reviewed the manuscript.

## NOTES

The authors declare the following competing financial interests: this work was supported by Eurofins DiscoverX, LLC, Eurofins DiscoverX Products, LLC and Eurofins Panlabs Discovery Services Taiwan, Ltd. M.K., J.K.W.D., L.M.G.L., G.P., N.B.S. and J.A.B are employees of Eurofins DiscoverX, LLC. R.M.H, J.L.J., J.E.L. and C.T.Y. are employees of Eurofins DiscoverX Products, LLC. T.Y.H., C.L.T. and C.Y.L. are employees of Eurofins Panlabs Discovery Services Taiwan, Ltd. The authors declare no other competing financial interest.

## REFERENCES

(1) Uprety, D., and Adjei, A. A. (2020) KRAS: From undruggable to a druggable Cancer Target. Cancer Treat. Rev. 89, 102070.

(2) Huang, L., Guo, Z., Wang, F., and Fu, L. (2021) KRAS mutation: from undruggable to druggable in cancer. Signal Transduct. Target. Ther. 6, 386.

(3) Malhotra, J., Nguyen, D., Tan, T., and Semeniuk Iii, G. B. (2024) Management of KRAS-mutated non-small cell lung cancer. Clin. Adv. Hematol. Oncol. 22, 67–75.

(4) Sahin, I. H., Saridogan, T., Ayasun, R., Syed, M. P., Gorantla, V., Malhotra, M., Thomas, R., Rhee, J., Zhang, J., Hsu, D., Singhi, A. D., and Saeed, A. (2024) Targeting KRAS oncogene for patients with colorectal cancer: A new step toward precision medicine. JCO Oncol. Pract. OP2300787.

(5) Stickler, S., Rath, B., and Hamilton, G. (2024) Targeting KRAS in pancreatic cancer. Oncol. Res. 32, 799–805.

(6) Ostrem, J. M., Peters, U., Sos, M. L., Wells, J. A., and Shokat, K. M. (2013) K-Ras(G12C) inhibitors allosterically control GTP affinity and effector interactions. Nature 503, 548–551.

(7) Bröker, J., Waterson, A. G., Smethurst, C., Kessler, D., Böttcher, J., Mayer, M., Gmaschitz, G., Phan, J., Little, A., Abbott, J. R., Sun, Q., Gmachl, M., Rudolph, D., Arnhof, H., Rumpel, K., Savarese, F., Gerstberger, T., Mischerikow, N., Treu, M., Herdeis, L., and W Fesik, S. (2022) Fragment optimization of reversible binding to the switch II pocket on KRAS leads to a potent, in vivo active KRASG12C inhibitor. J. Med. Chem. 65, 14614–14629.

(8) Patricelli, M. P., Janes, M. R., Li, L.-S., Hansen, R., Peters, U., Kessler, L. V., Chen, Y., Kucharski, J. M., Feng, J., Ely, T., Chen, J. H., Firdaus, S. J., Babbar, A., Ren, P., and Liu, Y. (2016) Selective Inhibition of Oncogenic KRAS Output with Small Molecules Targeting the Inactive State. Cancer Discov. 6, 316–329.

(9) Ma, X., Sloman, D. L., Duggal, R., Anderson, K. D., Ballard, J. E., Bharathan, I., Brynczka, C., Gathiaka, S., Henderson, T. J., Lyons, T. W., Miller, R., Munsell, E. V., Orth, P., Otte, R. D., Palani, A., Rankic, D. A., Robinson, M. R., Sather, A. C., Solban, N., Song, X. S., and Han, Y. (2024) Discovery of MK-1084: An Orally Bioavailable and Low-Dose KRASG12C Inhibitor. J. Med. Chem.

(10) Fell, J. B., Fischer, J. P., Baer, B. R., Blake, J. F., Bouhana, K., Briere, D. M., Brown, K. D., Burgess, L. E., Burns, A. C., Burkard, M. R., Chiang, H., Chicarelli, M. J., Cook, A. W., Gaudino, J. J., Hallin, J., Hanson, L., Hartley, D. P., Hicken, E. J., Hingorani, G. P., Hinklin, R. J., and Marx, M. A. (2020) Identification of the clinical development candidate MRTX849, a covalent KRASG12C inhibitor for the treatment of cancer. J. Med. Chem. 63, 6679–6693.

(11) Hallin, J., Engstrom, L. D., Hargis, L., Calinisan, A., Aranda, R., Briere, D. M., Sudhakar, N., Bowcut, V., Baer, B. R., Ballard, J. A., Burkard, M. R., Fell, J. B., Fischer, J. P., Vigers, G. P., Xue, Y., Gatto, S., Fernandez-Banet, J., Pavlicek, A., Velastagui, K., Chao, R. C., and Christensen, J. G. (2020) The KRASG12C Inhibitor MRTX849 Provides Insight toward Therapeutic Susceptibility of KRAS-Mutant Cancers in Mouse Models and Patients. Cancer Discov. 10, 54–71.

(12) Lanman, B. A., Allen, J. R., Allen, J. G., Amegadzie, A. K., Ashton, K. S., Booker, S. K., Chen, J. J., Chen, N., Frohn, M. J., Goodman, G., Kopecky, D. J., Liu, L., Lopez, P., Low, J. D., Ma, V., Minatti, A. E., Nguyen, T. T., Nishimura, N., Pickrell, A. J., Reed, A. B., and Cee, V. J. (2020) Discovery of a covalent inhibitor of KRASG12C (AMG 510) for the treatment of solid tumors. J. Med. Chem. 63, 52–65.

(13) Canon, J., Rex, K., Saiki, A. Y., Mohr, C., Cooke, K., Bagal, D., Gaida, K., Holt, T., Knutson, C. G., Koppada, N., Lanman, B. A., Werner, J., Rapaport, A. S., San Miguel, T., Ortiz, R., Osgood, T., Sun, J.-R., Zhu, X., McCarter, J. D., Volak, L. P., and Lipford, J. R. (2019) The clinical KRAS(G12C) inhibitor AMG 510 drives anti-tumour immunity. Nature 575, 217–223.

(14) Vasta, J. D., Peacock, D. M., Zheng, Q., Walker, J. A., Zhang, Z., Zimprich, C. A., Thomas, M. R., Beck, M. T., Binkowski, B. F., Corona, C. R., Robers, M. B., and Shokat, K. M. (2022) KRAS is vulnerable to reversible switch-II pocket engagement in cells. Nat. Chem. Biol. 18, 596–604.

(15) Wang, X., Allen, S., Blake, J. F., Bowcut, V., Briere, D. M., Calinisan, A., Dahlke, J. R., Fell, J. B., Fischer, J. P., Gunn, R. J., Hallin, J., Laguer, J., Lawson, J. D., Medwid, J., Newhouse, B., Nguyen, P., O’Leary, J. M., Olson, P., Pajk, S., Rahbaek, L., and Marx, M. A. (2022) Identification of MRTX1133, a noncovalent, potent, and selective KRASG12D inhibitor. J. Med. Chem. 65, 3123–3133.

(16) Peacock, D. M., Kelly, M. J. S., and Shokat, K. M. (2022) Probing the kras switch II groove by fluorine NMR spectroscopy. ACS Chem. Biol. 17, 2710–2715.

(17) Zhang, Z., Morstein, J., Ecker, A. K., Guiley, K. Z., and Shokat, K. M. (2022) Chemoselective Covalent Modification of K-Ras(G12R) with a Small Molecule Electrophile. J. Am. Chem. Soc. 144, 15916–15921.

(18) Zhang, Z., Gao, R., Hu, Q., Peacock, H., Peacock, D. M., Dai, S., Shokat, K. M., and Suga, H. (2020) GTP-State-Selective Cyclic Peptide Ligands of K-Ras(G12D) Block Its Interaction with Raf. ACS Cent. Sci. 6, 1753–1761.

(19) Yu, Z., He, X., Wang, R., Xu, X., Zhang, Z., Ding, K., Zhang, Z.-M., Tan, Y., and Li, Z. (2023) Simultaneous Covalent Modification of K-Ras(G12D) and K-Ras(G12C) with Tunable Oxirane Electrophiles. J. Am. Chem. Soc. 145, 20403–20411.

(20) Zheng, Q., Zhang, Z., Guiley, K. Z., and Shokat, K. M. (2024) Strain-release alkylation of Asp12 enables mutant selective targeting of K-Ras-G12D. Nat. Chem. Biol.

(21) Kessler, D., Gmachl, M., Mantoulidis, A., Martin, L. J., Zoephel, A., Mayer, M., Gollner, A., Covini, D., Fischer, S., Gerstberger, T., Gmaschitz, T., Goodwin, C., Greb, P., Häring, D., Hela, W., Hoffmann, J., Karolyi-Oezguer, J., Knesl, P., Kornigg, S., Koegl, M., and McConnell, D. B. (2019) Drugging an undruggable pocket on KRAS. Proc Natl Acad Sci USA 116, 15823–15829.

(22) Kim, D., Herdeis, L., Rudolph, D., Zhao, Y., Böttcher, J., Vides, A., Ayala-Santos, C. I., Pourfarjam, Y., Cuevas-Navarro, A., Xue, J. Y., Mantoulidis, A., Bröker, J., Wunberg, T., Schaaf, O., Popow, J., Wolkerstorfer, B., Kropatsch, K. G., Qu, R., de Stanchina, E., Sang, B., and Lito, P. (2023) Pan-KRAS inhibitor disables oncogenic signalling and tumour growth. Nature 619, 160–166.

(23) Zhang, Z., Guiley, K. Z., and Shokat, K. M. (2022) Chemical acylation of an acquired serine suppresses oncogenic signaling of K-Ras(G12S). Nat. Chem. Biol. 18, 1177–1183.

(24) Kirschner, T., Müller, M. P., and Rauh, D. (2024) Targeting KRAS diversity: covalent modulation of G12X and beyond in cancer therapy. J. Med. Chem. 67, 6044–6051.

(25) Bond, M. J., Chu, L., Nalawansha, D. A., Li, K., and Crews, C. M. (2020) Targeted Degradation of Oncogenic KRASG12C by VHL-Recruiting PROTACs. ACS Cent. Sci. 6, 1367– 1375.

(26) Galdeano, C., Gadd, M. S., Soares, P., Scaffidi, S., Van Molle, I., Birced, I., Hewitt, S., Dias, D. M., and Ciulli, A. (2014) Structure-guided design and optimization of small molecules targeting the protein-protein interaction between the von Hippel-Lindau (VHL) E3 ubiquitin ligase and the hypoxia inducible factor (HIF) alpha subunit with in vitro nanomolar affinities. J. Med. Chem. 57, 8657–8663.

(27) Li, L., Wu, Y., Yang, Z., Xu, C., Zhao, H., Liu, J., Chen, J., and Chen, J. (2021) Discovery of KRas G12C-IN-3 and Pomalidomide-based PROTACs as degraders of endogenous KRAS G12C with potent anticancer activity. Bioorg. Chem. 117, 105447.

(28) Hallin, J., Bowcut, V., Calinisan, A., Briere, D. M., Hargis, L., Engstrom, L. D., Laguer, J., Medwid, J., Vanderpool, D., Lifset, E., Trinh, D., Hoffman, N., Wang, X., David Lawson, J., Gunn, R. J., Smith, C. R., Thomas, N. C., Martinson, M., Bergstrom, A., Sullivan, F., and Christensen, J. G. (2022) Anti-tumor efficacy of a potent and selective non-covalent KRASG12D inhibitor. Nat. Med. 28, 2171–2182.

(29) Kemp, S. B., Cheng, N., Markosyan, N., Sor, R., Kim, I.-K., Hallin, J., Shoush, J., Quinones, L., Brown, N. V., Bassett, J. B., Joshi, N., Yuan, S., Smith, M., Vostrejs, W. P., Perez-Vale, K. Z., Kahn, B., Mo, F., Donahue, T. R., Radu, C. G., Clendenin, C., and Stanger, B. Z. (2023) Efficacy of a Small-Molecule Inhibitor of KrasG12D in Immunocompetent Models of Pancreatic Cancer. Cancer Discov. 13, 298–311.

(30) Fabian, M. A., Biggs, W. H., Treiber, D. K., Atteridge, C. E., Azimioara, M. D., Benedetti, M. G., Carter, T. A., Ciceri, P., Edeen, P. T., Floyd, M., Ford, J. M., Galvin, M., Gerlach, J. L., Grotzfeld, R. M., Herrgard, S., Insko, D. E., Insko, M. A., Lai, A. G., Lélias, J.-M., Mehta, S. A., and Lockhart, D. J. (2005) A small molecule-kinase interaction map for clinical kinase inhibitors. Nat. Biotechnol. 23, 329–336.

(31) Karaman, M. W., Herrgard, S., Treiber, D. K., Gallant, P., Atteridge, C. E., Campbell, B. T., Chan, K. W., Ciceri, P., Davis, M. I., Edeen, P. T., Faraoni, R., Floyd, M., Hunt, J. P., Lockhart, D. J., Milanov, Z. V., Morrison, M. J., Pallares, G., Patel, H. K., Pritchard, S., Wodicka, L. M., and Zarrinkar, P. P. (2008) A quantitative analysis of kinase inhibitor selectivity. Nat. Biotechnol. 26, 127–132.

(32) Davis, M. I., Hunt, J. P., Herrgard, S., Ciceri, P., Wodicka, L. M., Pallares, G., Hocker, M., Treiber, D. K., and Zarrinkar, P. P. (2011) Comprehensive analysis of kinase inhibitor selectivity. Nat. Biotechnol. 29, 1046–1051.

(33) Wodicka, L. M., Ciceri, P., Davis, M. I., Hunt, J. P., Floyd, M., Salerno, S., Hua, X. H., Ford, J. M., Armstrong, R. C., Zarrinkar, P. P., and Treiber, D. K. (2010) Activation state-dependent binding of small molecule kinase inhibitors: structural insights from biochemistry. Chem. Biol. 17, 1241–1249.

(34) Zhao, L., Zhao, J., Zhong, K., Tong, A., and Jia, D. (2022) Targeted protein degradation: mechanisms, strategies and application. Signal Transduct. Target. Ther. 7, 113.

(35) Zengerle, M., Chan, K.-H., and Ciulli, A. (2015) Selective small molecule induced degradation of the BET bromodomain protein BRD4. ACS Chem. Biol. 10, 1770–1777.

(36) Sakamoto, K. M., Kim, K. B., Verma, R., Ransick, A., Stein, B., Crews, C. M., and Deshaies, R. J. (2003) Development of Protacs to target cancer-promoting proteins for ubiquitination and degradation. Mol. Cell. Proteomics 2, 1350–1358.

(37) Nabet, B., Ferguson, F. M., Seong, B. K. A., Kuljanin, M., Leggett, A. L., Mohardt, M. L., Robichaud, A., Conway, A. S., Buckley, D. L., Mancias, J. D., Bradner, J. E., Stegmaier, K., and Gray, N. S. (2020) Rapid and direct control of target protein levels with VHL-recruiting dTAG molecules. Nat. Commun. 11, 4687.

(38) Sakamoto, K., Kamada, Y., Sameshima, T., Yaguchi, M., Niida, A., Sasaki, S., Miwa, M., Ohkubo, S., Sakamoto, J.-I., Kamaura, M., Cho, N., and Tani, A. (2017) K-Ras(G12D)-selective inhibitory peptides generated by random peptide T7 phage display technology. Biochem. Biophys. Res. Commun. 484, 605–611.

(39) Garrigou, M., Sauvagnat, B., Duggal, R., Boo, N., Gopal, P., Johnston, J. M., Partridge, A., Sawyer, T., Biswas, K., and Boyer, N. (2022) Accelerated Identification of Cell Active KRAS Inhibitory Macrocyclic Peptides using Mixture Libraries and Automated Ligand Identification System (ALIS) Technology. J. Med. Chem. 65, 8961–8974.

(40) Ergin, E., Dogan, A., Parmaksiz, M., Elçin, A. E., and Elçin, Y. M. (2016) Time-Resolved Fluorescence Resonance Energy Transfer [TR-FRET] Assays for Biochemical Processes. Curr. Pharm. Biotechnol. 17, 1222–1230.

(41) Olson, K. R., and Eglen, R. M. (2007) Beta galactosidase complementation: a cell-based luminescent assay platform for drug discovery. Assay Drug Dev. Technol. 5, 137–144.

(42) Nakane, A., Gotoh, Y., Ichihara, J., and Nagata, H. (2015) New screening strategy and analysis for identification of allosteric modulators for glucagon-like peptide-1 receptor using GLP-1 (9-36) amide. Anal. Biochem. 491, 23–30.

(43) Bonazzi, S., d’Hennezel, E., Beckwith, R. E. J., Xu, L., Fazal, A., Magracheva, A., Ramesh, R., Cernijenko, A., Antonakos, B., Bhang, H.-E. C., Caro, R. G., Cobb, J. S., Ornelas, E., Ma, X., Wartchow, C. A., Clifton, M. C., Forseth, R. R., Fortnam, B. H., Lu, H., Csibi, A., and Solomon, J. M. (2023) Discovery and characterization of a selective IKZF2 glue degrader for cancer immunotherapy. Cell Chem. Biol. 30, 235–247.e12.

(44) Wang, T., Li, Z., Cvijic, M. E., Krause, C., Zhang, L., and Sum, C. S. (2004) Measurement of β-Arrestin Recruitment for GPCR Targets, in Assay Guidance Manual (Sittampalam, G. S., Coussens, N. P., Brimacombe, K., Grossman, A., Arkin, M., Auld, D., Austin, C., Baell, J., Bejcek, B., Chung, T. D. Y., Dahlin, J. L., Devanaryan, V., Foley, T. L., Glicksman, M., Hall, M. D., Hass, J. V., Inglese, J., Iversen, P. W., Kahl, S. D., Kales, S. C., Lal-Nag, M., Li, Z., McGee, J., McManus, O., Riss, T., Trask, O. J., Weidner, J. R., Xia, M., and Xu, X., Eds.). Eli Lilly & Company and the National Center for Advancing Translational Sciences, Bethesda (MD).

(45) Swatek, K. N., and Komander, D. (2016) Ubiquitin modifications. Cell Res. 26, 399–422.

(46) Romero, C., Lambert, L. J., Sheffler, D. J., De Backer, L. J. S., Raveendra-Panickar, D., Celeridad, M., Grotegut, S., Rodiles, S., Holleran, J., Sergienko, E., Pasquale, E. B., Cosford, N. D. P., and Tautz, L. (2020) A cellular target engagement assay for the characterization of SHP2 (PTPN11) phosphatase inhibitors. J. Biol. Chem. 295, 2601–2613.

(47) McNulty, D. E., Bonnette, W. G., Qi, H., Wang, L., Ho, T. F., Waszkiewicz, A., Kallal, L. A., Nagarajan, R. P., Stern, M., Quinn, A. M., Creasy, C. L., Su, D.-S., Graves, A. P., Annan, R. S., Sweitzer, S. M., and Holbert, M. A. (2018) A High-Throughput Dose-Response Cellular Thermal Shift Assay for Rapid Screening of Drug Target Engagement in Living Cells, Exemplified Using SMYD3 and IDO1. SLAS Discov. 23, 34–46.

(48) Müller, C. W., Rey, F. A., Sodeoka, M., Verdine, G. L., and Harrison, S. C. (1995) Structure of the NF-kappa B p50 homodimer bound to DNA. Nature 373, 311–317.

(49) Schulze, J., Moosmayer, D., Weiske, J., Fernández-Montalván, A., Herbst, C., Jung, M., Haendler, B., and Bader, B. (2015) Cell-based protein stabilization assays for the detection of interactions between small-molecule inhibitors and BRD4. J. Biomol. Screen. 20, 180–189.

(50) Hong, D. S., Fakih, M. G., Strickler, J. H., Desai, J., Durm, G. A., Shapiro, G. I., Falchook, G. S., Price, T. J., Sacher, A., Denlinger, C. S., Bang, Y.-J., Dy, G. K., Krauss, J. C., Kuboki, Y., Kuo, J. C., Coveler, A. L., Park, K., Kim, T. W., Barlesi, F., Munster, P. N., and Li, B. T. (2020) KRASG12C Inhibition with Sotorasib in Advanced Solid Tumors. N. Engl. J. Med. 383, 1207– 1217.

(51) Awad, M. M., Liu, S., Rybkin, I. I., Arbour, K. C., Dilly, J., Zhu, V. W., Johnson, M. L., Heist, R. S., Patil, T., Riely, G. J., Jacobson, J. O., Yang, X., Persky, N. S., Root, D. E., Lowder, K. E., Feng, H., Zhang, S. S., Haigis, K. M., Hung, Y. P., Sholl, L. M., and Aguirre, A. J. (2021) Acquired resistance to KRASG12C inhibition in cancer. N. Engl. J. Med. 384, 2382–2393.

(52) Ash, L. J., Busia-Bourdain, O., Okpattah, D., Kamel, A., Liberchuk, A., and Wolfe, A. L. (2024) KRAS: biology, inhibition, and mechanisms of inhibitor resistance. Curr. Oncol. 31, 2024–2046.

(53) Tammaccaro, S. L., Prigent, P., Le Bail, J.-C., Dos-Santos, O., Dassencourt, L., Eskandar, M., Buzy, A., Venier, O., Guillemot, J.-C., Veeranagouda, Y., Didier, M., Spanakis, E., Kanno, T., Cesaroni, M., Mathieu, S., Canard, L., Casse, A., Windenberger, F., Calvet, L., Noblet, L., and Valtingojer, I. (2023) TEAD Inhibitors Sensitize KRASG12C Inhibitors via Dual Cell Cycle Arrest in KRASG12C-Mutant NSCLC. Pharmaceuticals (Basel) 16.

(54) Hagenbeek, T. J., Zbieg, J. R., Hafner, M., Mroue, R., Lacap, J. A., Sodir, N. M., Noland, C. L., Afghani, S., Kishore, A., Bhat, K. P., Yao, X., Schmidt, S., Clausen, S., Steffek, M., Lee, W., Beroza, P., Martin, S., Lin, E., Fong, R., Di Lello, P., and Dey, A. (2023) An allosteric pan-TEAD inhibitor blocks oncogenic YAP/TAZ signaling and overcomes KRAS G12C inhibitor resistance. Nat. Cancer 4, 812–828.

(55) Edwards, A. C., Stalnecker, C. A., Jean Morales, A., Taylor, K. E., Klomp, J. E., Klomp, J. A., Waters, A. M., Sudhakar, N., Hallin, J., Tang, T. T., Olson, P., Post, L., Christensen, J. G., Cox, A. D., and Der, C. J. (2023) TEAD Inhibition Overcomes YAP1/TAZ-Driven Primary and Acquired Resistance to KRASG12C Inhibitors. Cancer Res. 83, 4112–4129.

(56) Chapeau, E. A., Sansregret, L., Galli, G. G., Chène, P., Wartmann, M., Mourikis, T. P., Jaaks, P., Baltschukat, S., Barbosa, I. A. M., Bauer, D., Brachmann, S. M., Delaunay, C., Estadieu, C., Faris, J. E., Furet, P., Harlfinger, S., Hueber, A., Jiménez Núñez, E., Kodack, D. P., Mandon, E., and Schmelzle, T. (2024) Direct and selective pharmacological disruption of the YAP-TEAD interface by IAG933 inhibits Hippo-dependent and RAS-MAPK-altered cancers. Nat. Cancer.

